# Spatio-temporal mRNA dynamics in the early zebrafish embryo

**DOI:** 10.1101/2020.11.19.389809

**Authors:** Karoline Holler, Anika Neuschulz, Philipp Drewe-Boß, Janita Mintcheva, Bastiaan Spanjaard, Roberto Arsiè, Uwe Ohler, Markus Landthaler, Jan Philipp Junker

**Affiliations:** Max Delbrück Center for Molecular Medicine, Berlin Institute for Medical Systems Biology; Department of Biology, Humboldt University, Berlin, Germany; IRI Life Science, Institute of Biology, Humboldt Universität zu Berlin

**Keywords:** RNA localization, spatially-resolved transcriptomics, RNA labeling, zebrafish, early development, gastrulation, maternal factors

## Abstract

Early stages of embryogenesis depend heavily on subcellular localization and transport of maternally deposited mRNA. However, systematic analysis of these processes is currently hindered by a lack of spatio-temporal information in single-cell RNA sequencing. Here, we combined spatially-resolved transcriptomics and single-cell RNA labeling to study the spatio-temporal dynamics of the transcriptome during the first few hours of zebrafish development. We measured spatial localization of mRNA molecules with sub-single-cell resolution at the one-cell stage, which allowed us to identify a class of mRNAs that are specifically localized at an extraembryonic position, the vegetal pole. Furthermore, we established a method for high-throughput single-cell RNA labeling in early zebrafish embryos, which enabled us to follow the fate of individual maternal transcripts until gastrulation. This approach revealed that many localized transcripts are specifically transported to the primordial germ cells. Finally, we acquired spatial transcriptomes of two xenopus species, and we compared evolutionary conservation of localized genes as well as enriched sequence motifs. In summary, we established sub-single-cell spatial transcriptomics and single-cell RNA labeling to reveal principles of mRNA localization in early vertebrate development.

## Introduction

During embryonic development, initially pluripotent cells differentiate into a multitude of different cell types with distinct gene expression programs and spatial organisation. Advances in single-cell RNA sequencing (scRNA-seq)^1–4^ have made it possible to generate large single-cell atlases describing complex biological processes, including embryonic development of selected organisms. If the number of cells that are sampled is high enough, even extremely transient, and hence rare, states can be detected. This allows for an ordering of cells along an inferred pseudo-temporal trajectory^5–8^. Some of these approaches have been used successfully to reconstruct the cellular differentiation trajectories that underlie embryonic development in different species^9–12^. However, they fail to give insight into the earliest stages of embryonic development, which in many species are heavily regulated by RNA transport and intracellular localization of maternal transcripts^13,14^. In scRNA-seq, spatial information is lost and transcriptomic changes within a cell are not captured. In zebrafish for instance, which develop their body plan and all major organ primordia within 24h post fertilization (hpf), scRNA-seq does not resolve early patterning events that take place in the first 4 hours of development^9,10^.

While many methods for spatially-resolved transcriptomics have emerged in recent years^15,16^, state-of-the-art spatial RNA-seq methods typically have not reached the single-cell level yet^17^. Microscopy-based approaches using sequential fluorescent in situ hybridization hold great promise for spatial transcriptomics with sub-single-cell resolution^18–20^, but application of these methods to early embryos is technically challenging. Similarly, methods based on proximity labeling, which are powerful approaches for determining the transcriptome associated with different cellular compartments, require specific markers, transgenic engineering and have not been successfully applied to early vertebrate embryos yet^21,22^.

The temporal aspect of RNA expression is, by nature of the experiment, even harder to catch in single cells. While live microscopy based on fluorescent reporters is well established, methods for live measurement of transcript abundance typically consider only a couple of genes and are difficult to apply in live multicellular animals^23^. However, a cell’s ‘future transcriptome’ can, within certain limits, be inferred from RNA sequencing data by counting the occurrence of intronic reads^24^. Moreover, recent methods have made considerable progress in directly measuring the transcriptional history of single cells in cell culture by introducing modified nucleotides into newly synthesized RNA^25−29^.

In our study, we used a combination of spatially-resolved transcriptomics and RNA labeling to study the spatio-temporal dynamics of the transcriptome during the first few hours of zebrafish development. Specifically, we improved the tomo-seq method^30,31^ to measure RNA localization in one-cell stage zebrafish embryos with high spatial resolution. We used this information to systematically identify genes with sub-cellular localization patterns. Furthermore, we developed a protocol for single-cell RNA labeling in early zebrafish embryos that is compatible with high-throughput droplet microfluidics. This approach enabled us to follow the fate of individual maternal transcripts until gastrulation, and thereby deduce the biological function of the localized genes in embryonic development. We additionally investigated mRNA localization in an evolutionarily related system, oocytes from *Xenopus laevis* and *tropicalis*. This data allowed us to derive principles of mRNA localization in vertebrate oocytes, as well as evolutionary conservation and enriched sequence motifs.

## Results

For a systematic investigation of spatial RNA gradients in the zebrafish one-cell stage embryo, we established an enhanced, more sensitive version of the tomo-seq method^30^ (Methods): We embedded and oriented individual embryos at the one-cell stage (~30 min after fertilization) along the microscopically visible animal-vegetal axis. We then sectioned the cell and the yolk sac into 96 sections (Fig. 1a) and followed the tomo-seq protocol (Methods) for a total of three independent samples. We found that the majority of the mRNA is located in the blastodisc, which is positioned adherent to the yolk sac at the animal pole of the embryo (Fig. 1b, Fig. S1). To account for this pattern, we normalized transcript counts by total UMI counts per section, and recovered known localization patterns, as shown for important patterning genes like *dazl*, *trim36*, *grip2a*, *wnt8a* and *celf1* (Fig. 1c). Importantly, we found that our sub-single-cell tomo-seq library has high complexity, which enabled us to confidently determine spatial expression patterns of a large amount of genes: We found an average of 13.4 M unique transcripts (UMIs) per sample, and we observed that at the chosen sequencing depth (61 M reads on average), we are still far from reaching saturation, as determined by comparing UMI counts to read counts (Fig. 1d). Gene expression of individual replicates correlates well (R = 0.99, Fig. 1e, Fig. S1).

**Figure 1:**
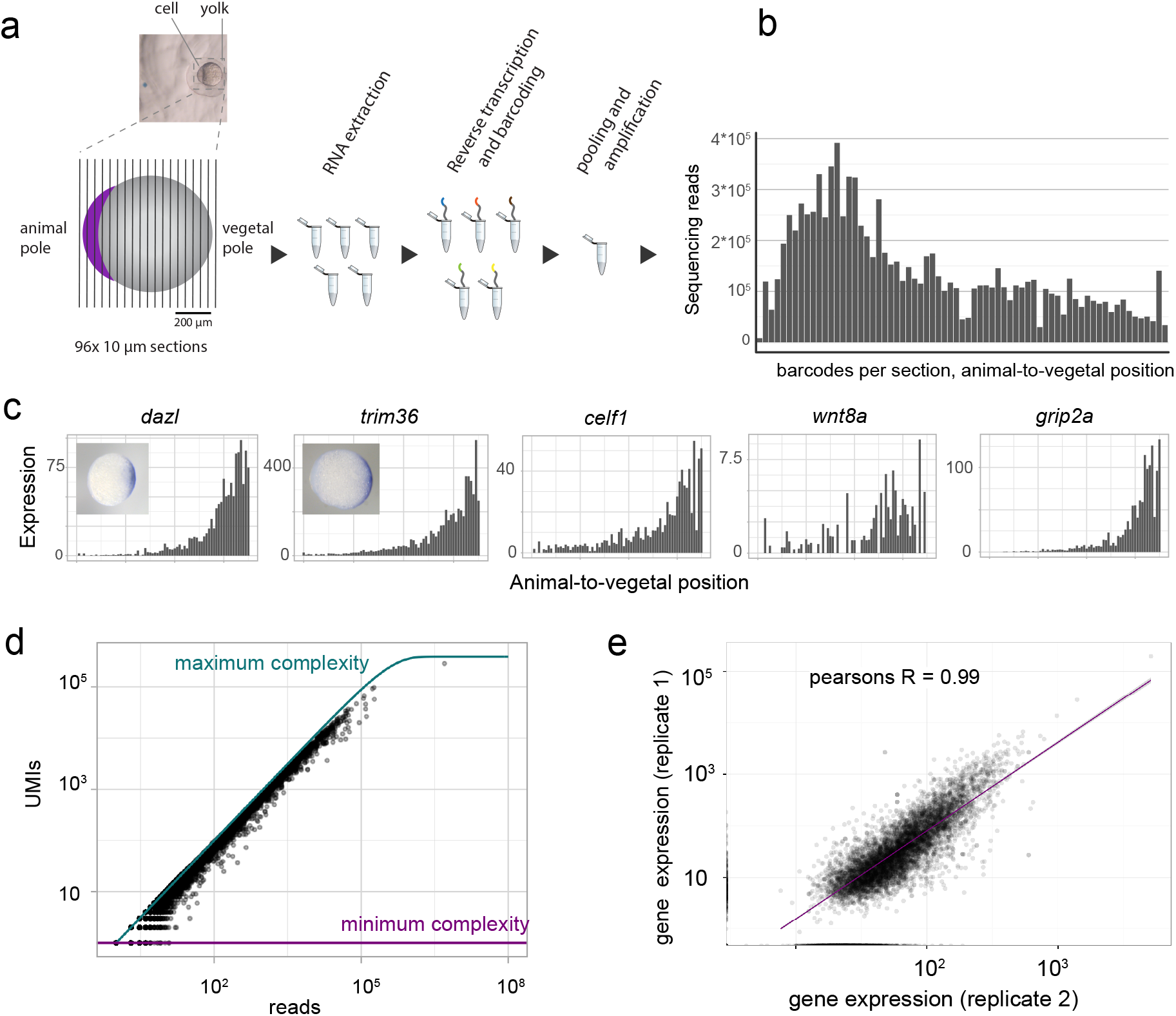
Low-input tomo-seq robustly recovers spatial mRNA gradients. a. Experimental outline: The embedded embryo is cryosectioned into 96 slices which are put into separate tubes. After adding spike-in control RNA, RNA is extracted. In a reverse transcription step, spatial barcodes are introduced. Samples are then pooled and amplified by invitro transcription and a final library PCR. Scale bars are 200 μm. b. Histogram shows raw transcript counts per section. c. Tomo-seq tracks for the known vegetally localized genes *dazl* and *trim36*, *celf1*, *wnt8a* and *grip2a* and whole-mount in-situ hybridizations (WISH) for *dazl* and *trim36*. d. Sequencing depth, shown as UMI saturation per gene. Maximum saturation is determined as by Grün *et al*. ^65^. e. Correlation of two tomo-seq experiments, line is a linear fit to the data.

In order to identify gene expression patterns in a systematic way, we clustered our spatial gene expression data based on a self-organizing map^30,32^, which sorted the cumulative gene expression traces along a linear axis of 50 profiles (supplementary table 1). As a result, we found three major groups of localized mRNA (Fig. 2a, S2): one localized to the animal side in profiles 1 to 8, one group of genes that was more or less equally distributed across all sections, and a third group of genes that was localized to the most vegetal part of the yolk sac in profiles 48-50. While the first group is likely an overlap between genes that had been localized to the animal pole before fertilization and transcripts that are transported by non-specific cell-directed cytoplasmic streaming upon fertilization^33^, the third group forms a distinct set of transcripts that were specifically transported and retained at the vegetal pole.

Since vegetally localized genes have been reported to play major roles in early development, especially in germ cell development and dorso-ventral axis specification^34,35^, and since this group of genes exhibited the most pronounced and reproducible spatial pattern in the one-cell stage embryo (Fig. S2), we decided to investigate it in more depth. We compared vegetally localized genes in profiles 48-50 between three replicates and found an overlap of 66 genes (Fig. 2b). A subset of the localized genes was not detected in one of the replicates (Fig. 2c), which was likely due to overall low expression of these genes. Another subset of genes that was defined as vegetally localized in one sample, was detected just below the threshold, in profiles 46 and 47, in another replicate (Fig. S2). Since manual inspection revealed that these genes had expression traces similar to genes previously annotated as vegetal (examples in Fig. S2), we decided to demand a vegetally localized gene to be in profiles 46-50 in all replicates, and in profile 48-50 in at least one of the replicates. With these criteria, we defined 97 genes to be localized vegetally, which increases the number of known vegetal genes by about tenfold. Moreover, this list includes all genes that to our knowledge have previously been shown to localize vegetally. We validated seven genes from this list, together with the animally localized gene *exd2*, by whole-mount in situ hybridization (WISH) (Fig. 2d). In summary, tomo-seq allowed us to determine subcellular RNA localization in the one-cell stage zebrafish embryo on the transcriptome wide level, which led to the identification of 97 genes that are specifically localized at the vegetal pole.

**Figure 2:**
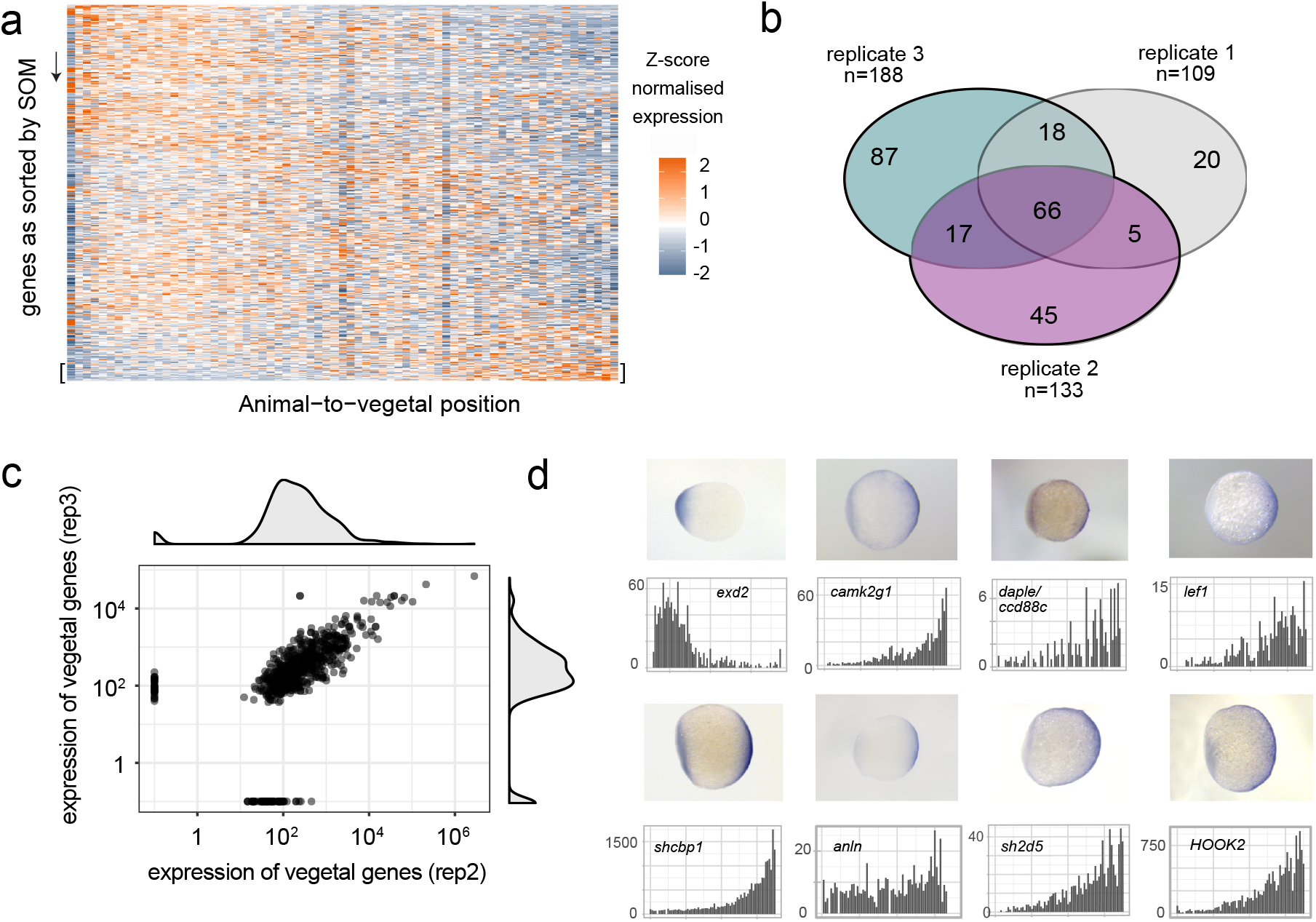
Tomo-seq of zebrafish one-cell stage embryos allows systematic identification of mRNA localization patterns. a. Heatmap of z-score normalized expression per section in a zebrafish one-cell stage embryo. Genes on the y-axis as sorted into profiles 1-50 by SOM, spatial position in the embryo on the x-axis. b. Vegetally localized genes per sample (profiles 48-50). c. Correlation of only vegetally localized genes of two replicates. Genes on the axes are only detected in one sample. d. Comparison of tomo-seq and WISH for selected newly described vegetally localized genes as well as the animally localized gene *exd2*.

To better understand the role of the vegetally localized genes in early development, it is important to follow the fate of maternal transcripts over time, in order to find out to which cell types they later contribute. The first major embryonic cell type decisions occur at gastrulation, which in the zebrafish happens at around 6 hpf^36^. Zygotic transcription starts at around 3 hpf, and gastrulation stages are characterized by a coexistence of maternal and zygotic transcripts. It is therefore crucial to distinguish maternal transcripts of localized genes from zygotic expression of the same genes.

We hence decided to develop an approach to distinguish maternal and zygotic transcripts transcriptome-wide and on the single cell level. Our method is based on single-cell RNA metabolic labeling (scSLAM-seq^25,36^), which enables us to distinguish maternal and zygotic transcripts by incorporation of the nucleotide analog 4-thiouridine (4sU). After a chemical derivatization step using iodoacetamide (IAA), labeled uridines are detected as T-to-C mutations upon sequencing^37^ (Fig. 3a). Several approaches for RNA labeling in single cells have been introduced recently^25–29,38^. However, these approaches are limited to cultured cells and have not been applied to live vertebrate embryos yet. Furthermore, they are mostly plate-based and (with the exception of Qiu et al.^29^) not compatible with high-throughput single-cell RNA-seq by droplet microfluidics. In order to study embryonic development, and to also capture rare cell types such as germ cells, it was crucial to overcome these limitations. We therefore developed a scSLAM-seq protocol that does not require cell lysis prior to IAA derivatization, which allowed us to load intact cells for droplet microfluidics scRNA-seq (Fig. 3a, Methods). To do so, cell membranes are permeabilized for IAA uptake by methanol fixation (Fig. S3). Compared to cultured cells, a major challenge in live emmbryos is to deliver the labeling reagent into the cells. Indeed, we found that addition of 4sU into the water did not yield high labeling efficiencies (Fig. S3). In bulk experiments, injection of 4-thiouridine-triphosphate (4sUTP) into one-cell stage zebrafish embryos has been used successfully for studying maternal-to-zygotic transcription^39^. Using the triphosphate 4sUTP has the additional advantage that the nucleotide analog is available immediately for incorporation into RNA without relying on further metabolic conversion. We observed efficient RNA labeling and successful conversion with IAA on bulk RNA upon 4sUTP injection (Fig. S3). We then proceeded to prepare single-cell suspensions at 50% gastrulation, fixed the cells with methanol and converted 4sUTP to a cytosine analogue in intact cells (Fig. 3a, Fig. S3). We then sequenced a total of 7472 cells with 10x Genomics Chromium, and analyzed the data with custom code (see Methods). Comparison of mutation rates in 4sUTP injected embryos with control samples confirmed that the T to C mutation rate is increased strongly and specifically (Fig. 3b). We found that the 4sUTP treatment resulted in a bimodal distribution of the T-to-C mutation frequency per gene (Fig. 3c), suggesting a good separation of labeled and unlabeled reads. The observed labeling efficiency of 5% corresponds to a low false negative rate of ~1% of unlabeled zygotic transcripts (Methods), which demonstrates that we can reliably distinguish maternal and zygotic transcripts.

**Figure 3:**
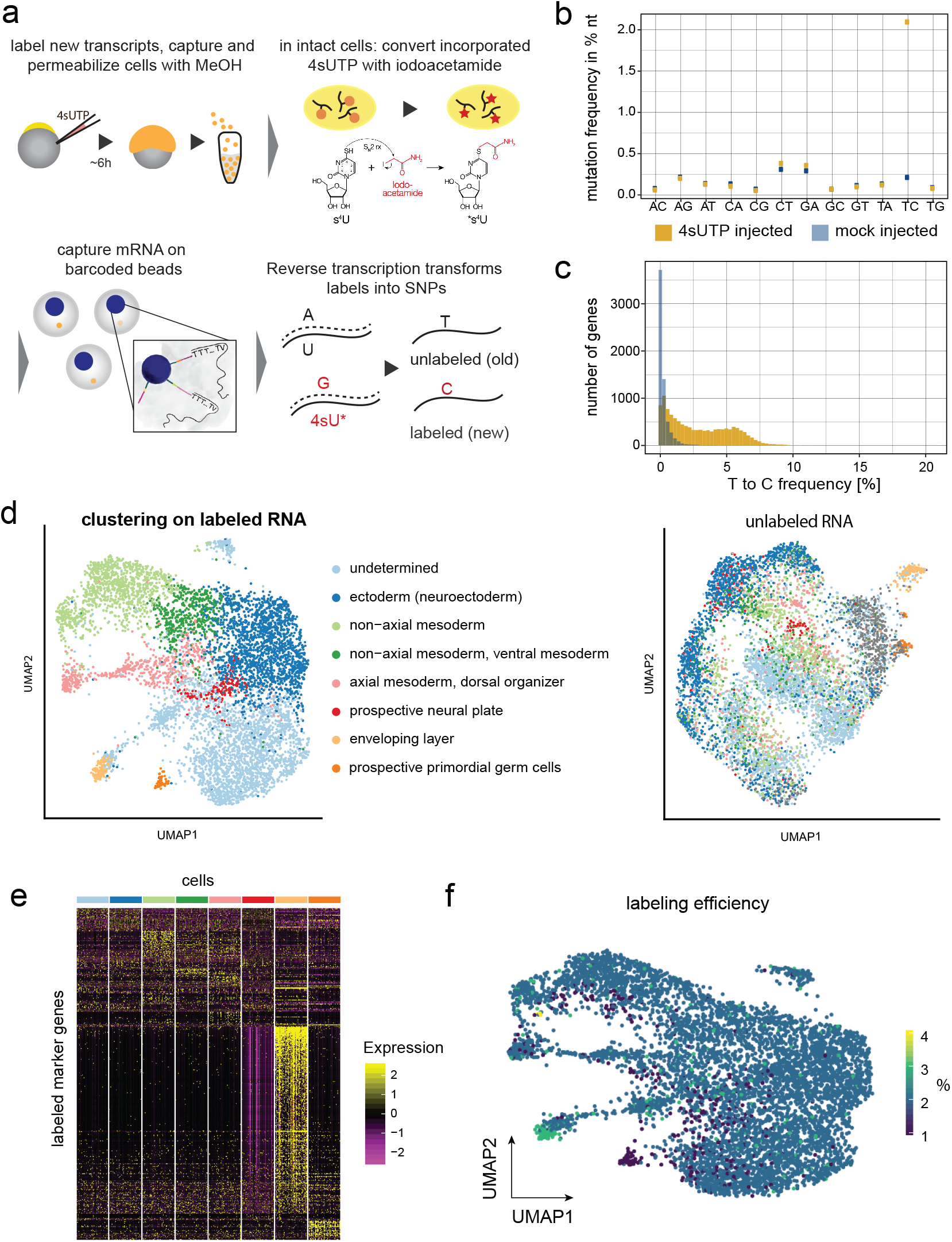
High-throughput single-cell RNA labeling in early zebrafish embryos. a. 4sUTP injection into zebrafish one-cell stage embryos, dechorionation, dissociation into single cells at gastrulation stage and methanol fixation (see Methods). Incorporated 4sU is converted in a SN2 reaction with iodoacetamide into a cytosine analogue. Single cell solution is then loaded onto a microfluidic device, chemical labels introduce SNPs during the first reverse transcription. b. Nucleotide mutation frequencies of a scSLAM-seq library after injecting 4sUTP or Tris and quality filtering of the data. c. Histogram of T-to-C mutations in 4sUTP- and Tris-injected embryos. d. UMAP representation of cells based on labeled RNA (left side) and unlabeled RNA (right side). For the latter, we imposed cell identities as determined on the basis of labeled RNA. e. Marker gene expression of labeled cells in different cell types. Cell number per cluster was downsampled to equal numbers. f. Transcript labeling efficiency in single cells in percent, projected on the UMAP representation for labeled RNA.

Unsupervised clustering of cells, using the information of the labeled mRNA, resulted in eight cell clusters (Fig. 3d) with defined marker gene expression (Fig. 3e, supplementary table 2). We then clustered cells based on their unlabeled mRNA (Fig. 3d) and imposed cell identities as defined based on labeled mRNA. As expected, clustering based on unlabeled (maternal) mRNA separated cell types much less than clustering on labeled (zygotic) RNA, with the notable exception of the enveloping layer and the primordial germ cells (PGCs). These two cell types had the most distinct marker gene signature (Fig. 3e, supplementary table 2), and the cells of the enveloping layer were characterized by a particularly high labeling rate (Fig. 3f), which indicates high transcriptional or proliferative activity. The PGCs, on the other hand, display the lowest labeling rates among all cells at this developmental stage (Fig. 3f), in agreement with reports that show very slow increase of prospective PGCs before gastrulation^40,41^.

Next, we set out to assess if any of the maternal, vegetally localized genes were over-represented in specific cell types. At 6 hpf, we still detected unlabeled RNA for 91 of the 97 genes that were localized at the one-cell stage. We filtered out lowly expressed genes, and for the remaining 47 genes we calculated the expression fold change for each of the cell types compared to all other cell types (Fig. 4a). We found that the vegetally localized genes were significantly enriched in PGCs (p-value = 4.67*10^−5^), with 28 of them being marker genes of that particular cluster (supplementary table 3). The logarithmic fold enrichment of vegetal genes in PGCs follows a bimodal distribution (Fig. 4b, dashed line), suggesting two subpopulations of vegetal genes. Indeed, we can deconvolve the bimodal distribution into two normal distributions, where one resembles the distribution of randomly sampled genes (Fig. 4b light blue and gray), while the other has a significantly higher mean fold change (Fig. 4b dark blue, p-value=1.7*10^−4^), suggesting a role of these genes in germ cell specification or development. We show the average expression at 6 hpf for some of the new candidates (*sh2d5*, *itpkca*, *ndel1b*, *anln*, *krtcap2* and *ppp1r3b*) in Figure 4c, next to the remaining maternal expression of well-established germ cell factors (Fig. 4d). Interestingly, the transcripts of vegetally localized genes with a known role in axis formation (*wnt8a* and *syntabulin*) cannot be detected any more in the maternal transcriptome, which suggests that such factors are degraded more rapidly than germ cell factors. In summary, our scSLAM-seq analysis revealed that a large number of the vegetally localized transcripts are later transported to primordial germ cells, thereby allowing us to identify a set of novel candidate genes with a potential function in germ cell specification and differentiation.

**Figure 4:**
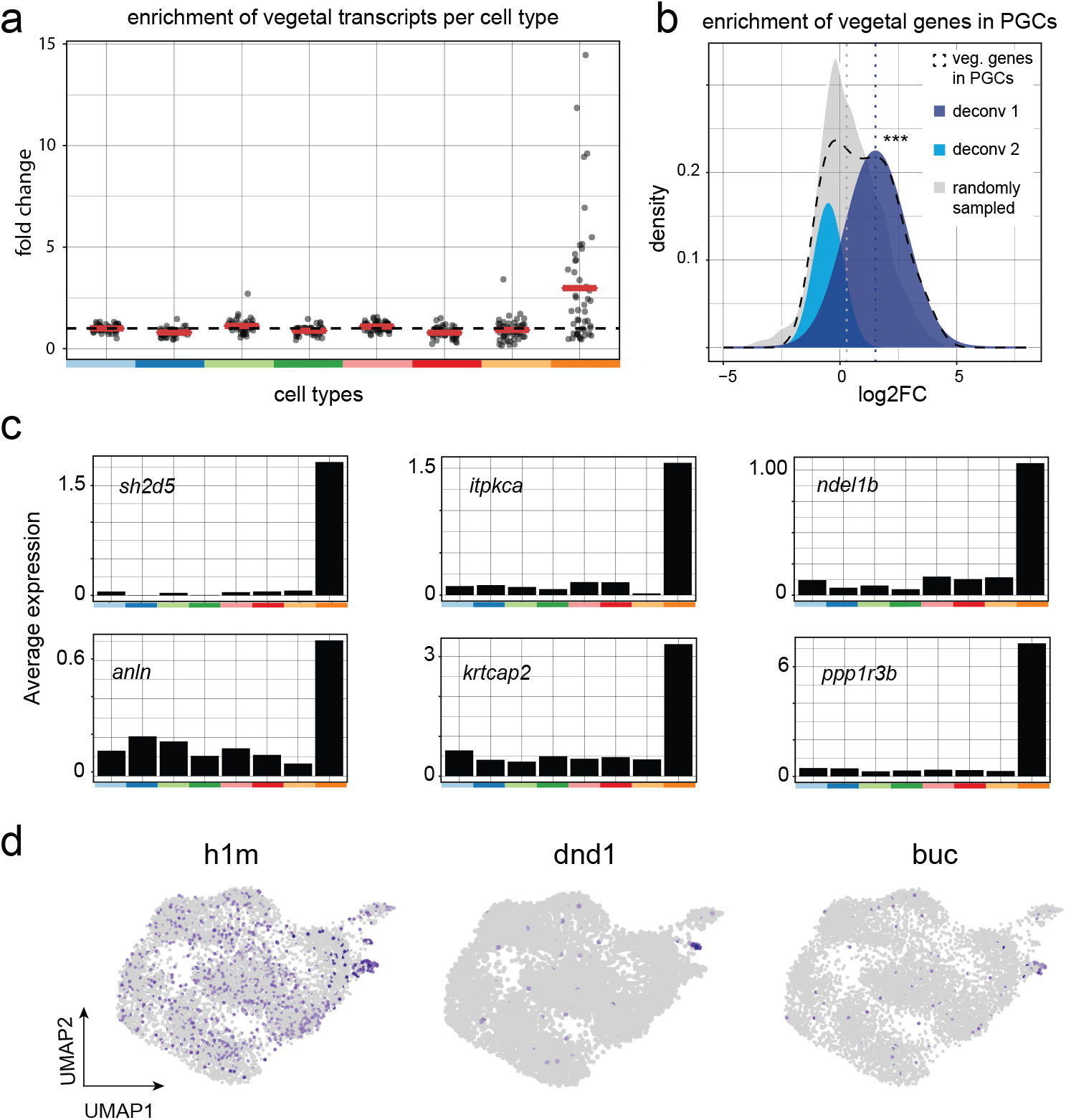
High-throughput scSLAM-seq follows the fate of maternal transcripts until gastrulation. a. Fold change enrichment of maternal vegetally localized genes for different cell types vs. all other cells. Genes with an average expression lower than 0.1 were excluded from this analysis. Red bars represent mean values. b. Deconvolution of the bimodal distribution of vegetally localised genes in PGCs (black dashed line) into two normal distributions (light and dark blue). Mean value of dark blue distribution is significantly higher than of a randomly sampled distribution (m_gray_ = 0.4, m_darkblue_ = 1.52, p-value = 1.7*10^−4^, Welchs t-test). c. Average expression of most highly enriched genes in PGCs in different cell types. d. Unlabeled RNA expression of established germ cell markers on a UMAP representation.

We next decided to determine the conservation of germ cell factors by comparing vegetally localized genes in zebrafish and xenopus. Our choice of xenopus was motivated by reports showing that 3’UTR sequences of a zebrafish germ plasm gene can drive transcript localization in frog oocytes^42^, and furthermore that the localization machineries of two different xenopus species, *X. tropicalis* and *X. laevis*, are functionally overlapping^42,43^, which suggests that RNA localization is driven by common cis-regulatory elements. Importantly, xenopus as well as zebrafish use the vegetal pole to store factors for germ cell specification and dorso-ventral axis determination, which additionally suggests functional similarity despite a considerable evolutionary distance (Fig. 5a). Since existing xenopus datasets are derived from pooled samples and do not provide a comparable spatial resolution^44–46^, we decided to produce tomo-seq datasets of mature oocytes from *X. tropicalis* and *X. laevis*, with two replicates for each species (10 μm resolution for *X. tropicalis*, 16 and 18 μm resolution for *X. laevis*).

**Figure 5:**
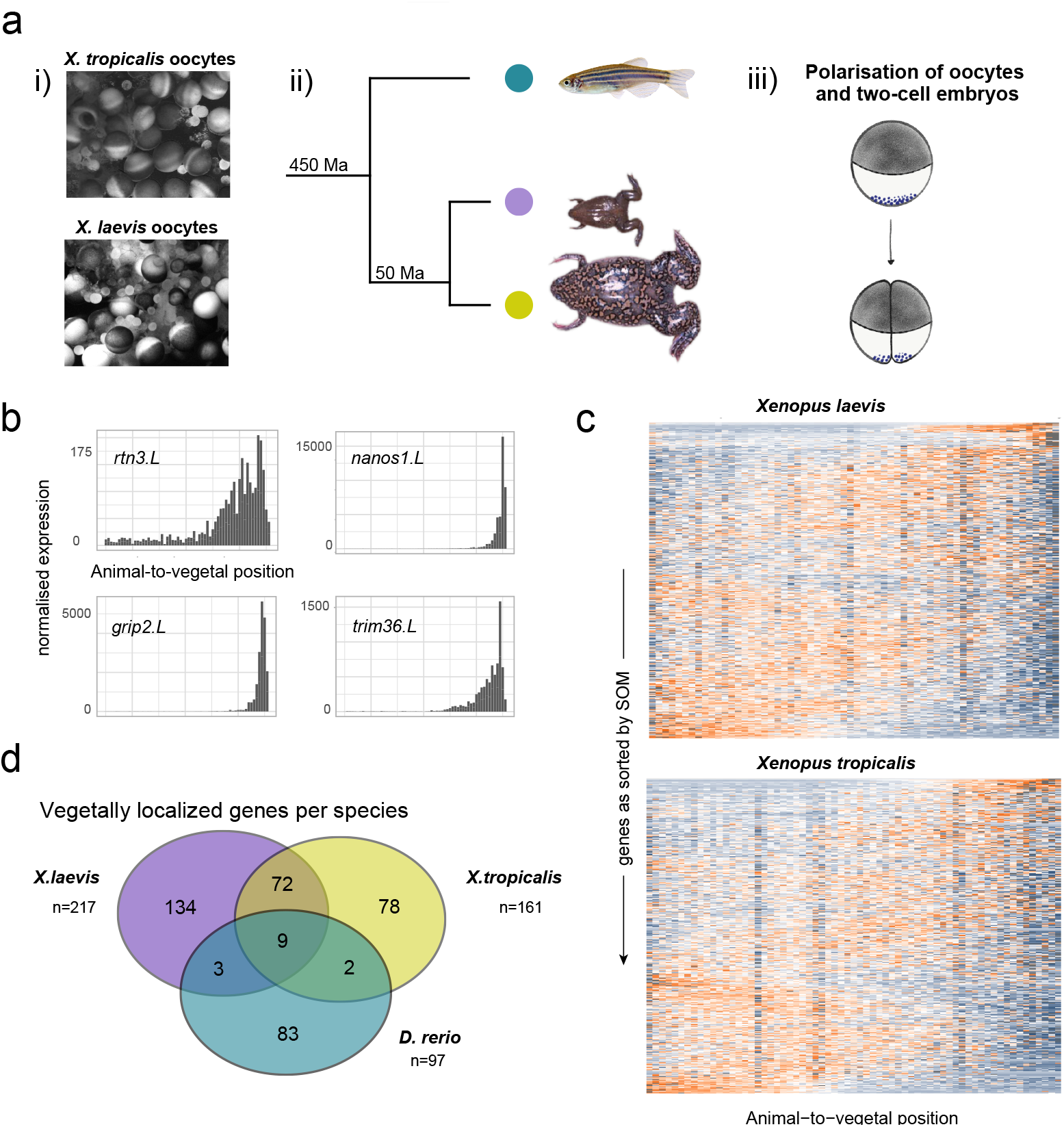
Evolutionary conservation of mRNA localization between zebrafish and xenopus. a. i) Light microscopy view of whole oocyte lobes from *X. laevis* and *X. tropicalis* before dissociation. ii) Phylogenetic distance of xenopus species and zebrafish. iii) Deposition of germ plasm and dorsal factors in xenopus oocytes and after first cell division. b. Tomo-seq tracks of vegetally localised genes *rtn3.L*, *nanos1.L*, *grip2.L* and *trim36.L* in *X. laevis*. c. Heatmap of z-score normalized expression per section in xenopus oocytes. Genes on the y-axis as sorted into profiles 1-50 by SOM, spatial position on the x-axis. d. Overlap of vegetally localized genes in zebrafish and xenopus species, considering only genes that were expressed in all three species at the respective developmental stage.

After excluding lowly expressed genes and normalizing to the same number of transcripts per section, we recovered known localization patterns for important developmental factors (Fig. 5b). As before, we calculated cumulative expression patterns and clustered them with self-organizing maps (Fig. 5c, S5, supplementary table 4, 5). In *X. tropicalis*, we found 151 genes to be localized animally (1.5%) and 161 to be localized vegetally (1.6%), for *X. laevis* we identified 245 genes localized to the animal pole (1.9%) and 216 genes to the vegetal pole (1.7%). In accordance with a previous study^44^, the interspecies overlap of localized genes was relatively low for these two closely related frog species – 30% for animally localized genes and 50% for vegetally localized genes. One important difference between high-resolution tomo-seq data and earlier studies of *X. laevis* and *X. tropicalis* is the identification of a distinct group of animally localized genes and their corresponding motifs (Fig. S5). The existence of animally localized genes was previously controversial, since either very few (0.2%, Owens et al.^45^) or a large majority of genes (94.4%, Sindelka et al.^46^) were found to be enriched at the animal pole. This highlights the advantages of our high-resolution analysis of subcellular RNA localization.

While the overlap of the vegetal genes between the two xenopus species with 81 genes was considerable, we only found nine genes to localize vegetally in all three species (Fig. 5d), showing a surprisingly variable transcript composition at the vegetal pole given the reported high degree of conservation of the localization machinery^42^. However, this analysis allowed us to propose that these nine genes, including known factors like *dazl* and *syntabulin*, but also less well characterized genes like *camk2g1* and *ppp1r3b*, have a conserved function in germ cell development or dorso-ventral axis development. *Camk2g1*, for instance, was found to be transported to the PGCs in our scSLAM-seq analysis (Fig. 4), which indicates a conserved role of this gene in germ cell specification. Interestingly, *anln* is PGC enriched and localized in zebrafish, and it is vegetally localized in *X. tropicalis* but not in *X. laevis*. The 3’UTR of *X. laevis anln* has a 1 kb long deletion, suggesting a functional contribution of that sequence to the localization (Fig. S5). Table 1 gives an overview of the nine genes with conserved localization, their described cellular function (Xenbase.org, zfin.org) and the protein class of the translated product.

**Table 1:**
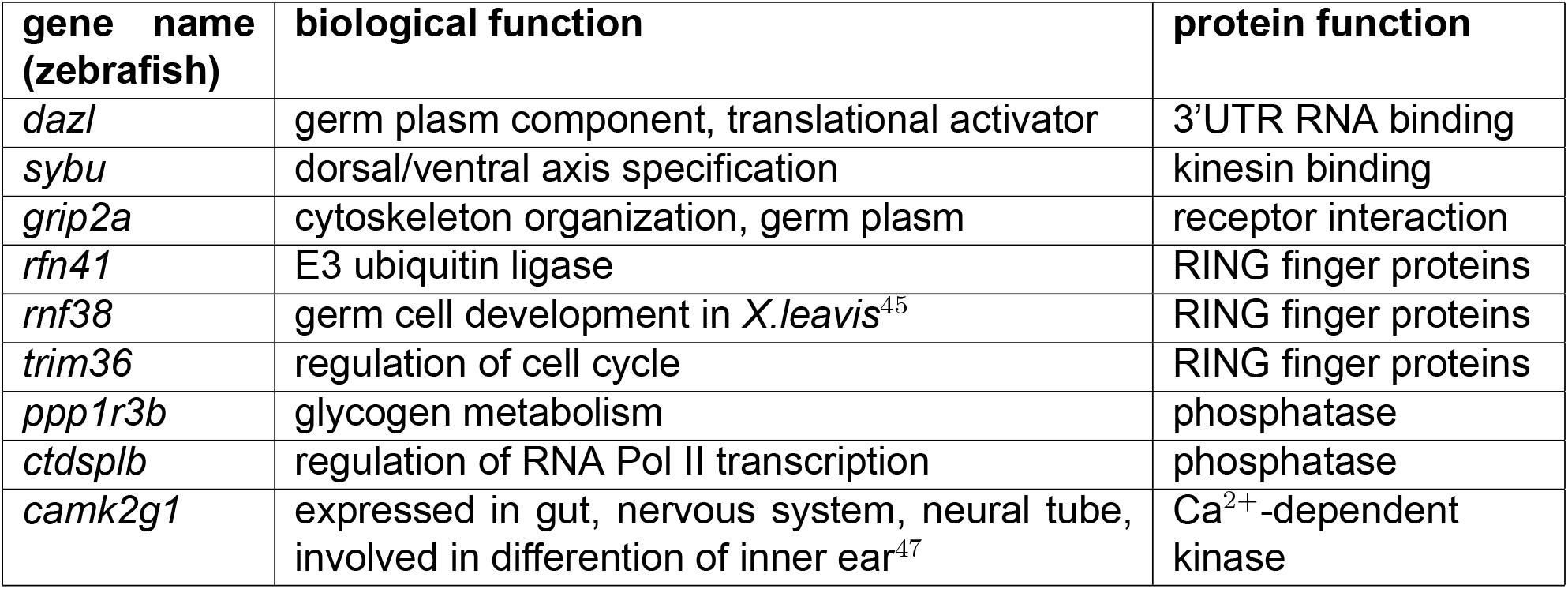
Genes with conserved vegetal localization in zebrafish, *X. tropicalis* and *X. laevis*.

Intracellular transcript localization is driven by cis-regulatory localization elements, present mainly in 3’ untranslated regions (UTRs) of RNA molecules^42,48–51^. However, the exact nature of the localization motifs in early embryos has largely remained elusive. We therefore reasoned that our transcriptome-wide datasets of mRNA localization in three species might now open the door towards a more systematic analysis of these sequence elements. To this end, we decided to investigate shared sequence features of vegetally localized genes. Since tomo-seq detects only 3’ ends of transcripts, we performed bulk RNA-seq of one-cell stage zebrafish embryos in order to computationally identify expressed isoforms^52^ (Methods). In total, we detected 216 expressed isoforms of vegetally localized genes in zebrafish. We found that the 3’UTR sequences of vegetally localizing genes are on average 1.7-fold longer than for the background (p-value < 2.2*10^−16^) (Fig. 6a). In contrast to this, we found only moderate differences in length of coding sequences (Fig. 6b) and expression level (Fig. 6c), and no differences in GC content of 3’ UTRs (Fig. S6). Longer 3’UTRs of vegetally localized genes could reflect complex cellular regulation of these transcripts in regard to localization and anchoring to the cytoskeleton, but could also be at least partially related to other regulatory processes such as translational activity and RNA stability^53^. Finally, we searched for common cis-regulatory motifs by performing a k-mer enrichment analysis^54^ of the 3’UTRs (Fig. 6d, Methods). We detected variations of a CAC core, several motifs containing a GUU sequence that has not been described yet, and a poly-U stretch that was previously linked to increased RNA stability^54,55^.

**Figure 6:**
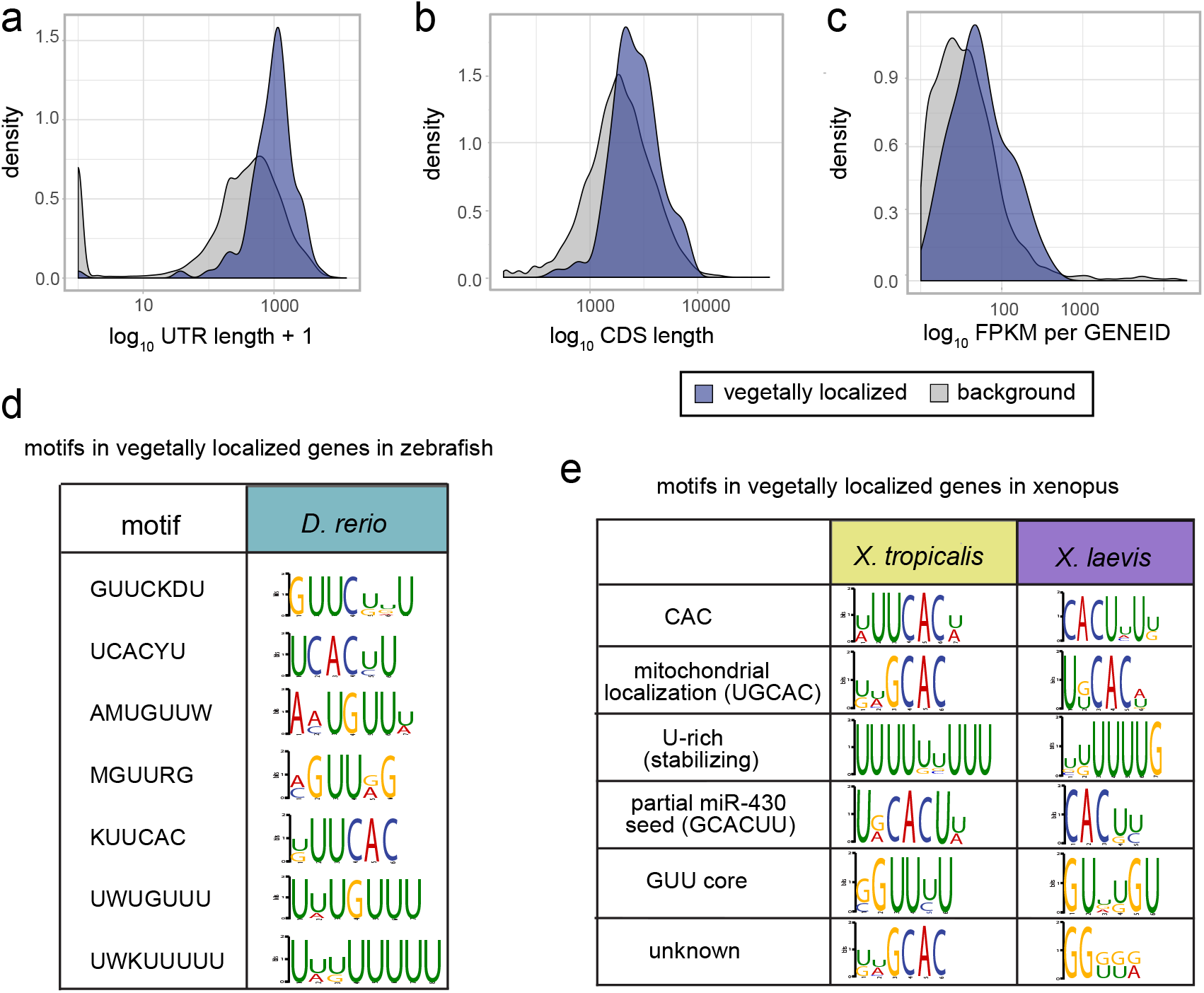
3’UTR characteristics of vegetally localized genes. a-c. Comparison of sequence characteristics of expressed isoforms of vegetally localized to all genes. a. Weighted 3’UTR lengths: Isoforms contribute according to their relative expression, mean(vegetal genes) = 1.06 kb, mean(background) = 0.6 kb, p-value < 2.2*10^−16^ (two-sample Wilcoxon test). b. Weighted lengths of coding regions, mean(vegetal genes) = 2.78 kb, mean(background) = 2.42 kb, p-value = 7.655*10^−14^ (two-sample Wilcoxon test). c. FPKM sum per gene ID, IDs with less than 10 FPKM were omitted. Mean expression of vegetal genes 64.1 FPKM, mean of background 37.4, p-value < 2.2*10^−16^, (two-sample Wilcoxon test). d. Results of the kmer enrichment analysis of 3’UTRs of 216 expressed isoforms, zebrafish vegetally localized genes. Top seven motifs and logos. e. Results of the kmer enrichment analysis of the longest 3’UTR of vegetally localized genes in *X. laevis* and *X. tropicalis*, top six motifs, and their respective description based on previous publications.

We next performed a k-mer enrichment analysis for the two xenopus species by using the longest annotated 3’UTR isoform (Fig. 6e, S6). In accordance with previous studies^48,56,57^, and similar to our results for zebrafish, we found an enrichment of CAC-containing motifs in vegetally localized genes. Interestingly, we found the same polyU-motif as in zebrafish data, suggesting a conserved role in stability of maternal RNA. In *X. tropicalis*, we also found a motif consisting of the same GUU core we identified in zebrafish (Fig. 6d); however, the respective local sequence environment differed. In summary, we found a relatively high conservation of 3’UTR sequence motifs, which contrasts with the rather low conservation of vegetally localized genes that we observed in Figure 5.

## Discussion

We here established improved versions of two methods, the tomo-seq approach for spatially-resolved transcriptomics, and single-cell SLAM-seq for RNA labeling. In tomo-seq, we achieved sub-single-cell resolution in zebrafish embryos at the one-cell stage. Importantly, we observed that the complexity of the tomo-seq libraries was not a limiting factor, suggesting that our approach may be applicable to even smaller samples containing less mRNA. The tomo-seq method is well suited for spatial transcriptomics in the one-cell stage zebrafish embryos, since we expect the most striking patterns along the animal-vegetal axis. However, in systems with more intricate spatial patterns, different approaches for spatially-resolved transcriptomics in 2D or 3D may be more suitable.

For scSLAM-seq, we achieved two important advances: We made the method compatible with high-throughput scRNA-seq based on widely-used droplet microfluidics approaches by performing the chemical derivatization of 4sU in intact methanol-fixed cells. Furthermore, we successfully labeled zygotically transcribed RNA in early zebrafish embryos by injecting 4sUTP into the zygote. scSLAM-seq is a universal approach for following the fate of RNA molecules over time, and we anticipate that this approach will emerge as a powerful method for short-term fate tracking of RNA molecules in living organisms. However, it is important to note that efficient delivery of 4sU into other live animals may require different approaches depending on the species and the organ system.

The combination of sub-single-cell tomo-seq and single-cell RNA labeling generates important synergy by allowing transcriptome-wide measurement of spatio-temporal RNA dynamics. We used this combination of techniques for a systems-level analysis of RNA dynamics in early zebrafish development, which gave us access to developmental events that are not captured by conventional scRNA-seq. Beyond this specific biological application, we anticipate that the combination of spatial transcriptomics and RNA labeling will find important applications for many other questions, such as tissue remodeling in disease conditions or analysis of cell-cell signaling interactions in vivo.

Besides the methodology presented here, another major output of this work consists in the transcriptome-wide resource of localized genes in three vertebrate species. While highresolution atlases of transcript localization have been established in Drosophila oocytes based on automated microscopy^58^, no comparable datasets exist for early vertebrate development, with the exception of low-resolution spatial analysis of xenopus oocytes^44–46^. Our analysis provides a shortlist of candidate genes with a potential role in early development, including genes like the phosphatase *ppp1r3b* or the kinase *camk2g1*, which have no known function in early embryogenesis but are vegetally localized in all three species. We observed that many vegetally localized transcripts are later specifically transported into the primordial germ cells, suggesting that specification of the PGCs is one of the main functions of the localized genes discovered here. Interestingly, we observed a relatively low conservation of localized genes, but a rather high conservation of enriched motifs in 3’UTRs. While it is possible that our analysis underestimates the true degree of conservation of vegetal localization due to the difficulty of reliably calling localization patterns for lowly expressed genes, this observation raises the question whether the function of genes involved in e.g. PGC specification is conserved, even if the localization pattern is not.

Asymmetric localization of mRNA molecules is a pervasive phenomenon in the animal kingdom^59−61^ and provides an important layer of gene regulation in a variety of different cell types by e.g. restricting translation spatially^59,62^ or by controlling translation efficiency^60^. While the exact nature of the localization motifs in early embryos have largely remained elusive, there are indications that secondary structures^63,64^ or sequence-dependent piRNA adhesion traps might be involved^65^. While our high-resolution spatial transcriptomics data allowed a systematic analysis of enriched k-mers, the results probably do not reveal the full mechanism, since we did not identify a single motif that explains localization of all genes. This, together with the observation that 3’UTRs of vegetally localized genes are longer than for other genes, suggests more complex and potentially longer regulatory elements than the k-mers analyzed here. We speculate that the combination of sub-single-cell tomo-seq with the injection of 3’UTR fragments may in the future provide further insights into the molecular mechanisms underlying RNA localization.

## Acknowledgements

The authors thank Jana Richter for her work on whole-mount in-situ hybridizations, Ronny Schäfer for zebrafish injections, and Nora Fresmann for her kind support with scSLAM-seq experiments during the Sars-Cov2 lockdown of the laboratory. We thank Moritz Ophaus who produced X. laevis tomo-seq libraries as a rotation student in the lab, and Roberto Moreno Ayala for support with cell type identification of single cell data. We also acknowledge support by MDC/BIMSB core facilities (zebrafish, genomics, bioinformatics). Work in J.P.J.’s lab was supported by a European Research Council Starting Grant (ERC-StG 715361 SPACEVAR) and a Helmholtz Incubator grant (Sparse2Big ZT-I-0007).

## Author contributions

K.H. performed zebrafish tomo-seq experiments and data analysis. Tomo-seq of *X. tropicalis*, analysis and comparison to *X. laevis* data was done by J.M. as part of her master thesis under supervision of K.H. and J.P.J.. Kmer enrichment analysis was done by P.B.. B.S. built the mapping pipeline, adapted self-organising maps to tomo-seq data and assisted with statistical data analysis. Experimental method development of scSLAM-seq was done by A.N. and R.A., and A.N. developed the computational pipeline for scSLAM-seq analysis. Single cell data was jointly analyzed by K.H and A.N., PGC specific and statistical analyses were performed by K.H. and B.S.. All authors discussed and interpreted the results. The paper was written by K.H. and J.P.J..

## Methods

### Animal methods

#### Breeding of zebrafish

Fish were maintained according to standard laboratory conditions. All animal procedures were conducted as approved by the local authorities (LAGeSo, Berlin, Germany). For embryo experiments, we set up group crosses of AB wild type fish.

#### Preparation of frog oocytes

Oocyte lobes were ordered from the European Xenopus Resource Centre (University of Portsmouth), manually dissected with forceps on agarose plates, and gently dissociated with liberase as described by Claussen *et al*^67^.

### Laboratory methods

#### Tomo-seq

Zebrafish embryos were harvested 20 min. after fertilization. Individual embryos were embedded in OCT medium under a dissection microscope and oriented along the animalvegetal axis with tungsten needles. Since the transparency of the embryo makes the embryo invisible after freezing the block, we marked the starting point for the blind collection of sections with a blue polyacrylamide bead (BIORAD). Before snap-freezing the cryomold on dry ice, we took a picture to calculate the distance between the edge of the block and the polyacrylamide bead in Fiji.

We sectioned the blocks into 96 sections (thickness 10 μm), followed by Trizol RNA extraction as described in Holler and Junker^31^. Pelleted RNA was directly dissolved in a mix of dNTPs and barcoded poly-dT primers, and was reverse-transcribed with SuperScript II. Primer design was inspired by CELseq2 (Hashimshony *et al*., 2016), using 8 nt barcodes, 6 nt UMIs, and a different adapter design.

The following steps include linear amplification with IVT, RNA fragmentation, 2nd reverse transcription and library PCR; as described in detail previously^31^.

For xenopus oocytes we used the same protocol, but adjusted the section thickness according to the sample diameter.

#### Bulk RNA sequencing

Embryos were harvested 20 minutes after fertilization and directly put into Trizol. We extracted RNA with chloroform and isopropanol, and dissolved the pelleted RNA in nucleasefree water. Quality of the RNA was checked on a bioanalyzer RNA pico chip. We then prepared full length sequencing libraries with the Illumina TruSeq stranded mRNA kit. The samples were sequenced on Illumina HiSeq4000.

#### WISH

Zebrafish embryos were fixed 20 min. after fertilization in 4% PFA for 2 h. Whole mount *in situ* hybridization was performed as in Thisse *et al*^68^.

#### scSLAM-seq

##### 4sUTP injections

We injected zebrafish embryos directly after fertilization with 4 nl 4sUTP (12.5 mM, SigmaAldrich, in 10 mM Tris•HCl pH 7.4, Carl Roth). At 50% epiboly we removed the chorions, then continued incubation until shield stage.

##### Cell fixation and iodoacetamide treatment

We dissociated 10 shield stage embryos per sample by gently pipetting up and down in deyolking buffer (55 mM NaCl, 1.8 mM KCl, 1.25 mM NaHCO3 in HBSS, Life Technologies). For cell fixation we added cold methanol (Carl Roth) until a final concentration of 80%. We then fixed the cells at −20°C for 30 min. For chemoconversion, we added 1 M iodoacetamide (Sigma-Aldrich) in 80% methanol and 20% HBSS to a final concentration of 10 mM, and gently agitated the mixture at room temperature, overnight, in the dark.

##### Rehydration and preparation for scRNA-seq

To inactivate the iodoacetamide, we spun down the cells at 1,000 g for 5 min and resuspended in quenching buffer (DBPS, Gibco, 0.1% BSA, Sigma-Aldrich, 1 U/μl RNaseOUT, Life Technologies, 100 mM DTT, Carl Roth) and incubated at room temperature for 5 min. After spinning down again, we resuspended them in DPBS containing 0.01% BSA, 0.5 U/μl RNaseOUT and 1 mM DTT. The cells were then passed through a 35 μm strainer, counted, and immediately loaded onto a 10x Chromium system using the 3’ kit (V2 and V3).

##### Library preparation and sequencing

We prepared sequencing libraries according to the manufacturer’s instructions and sequenced them on Illumina HiSeq4000 and NextSeq500 systems.

##### Dot blots for detection of incorporation and IAA derivatization of 4sUTP

We biotinylated extracted RNA using the following mixture: 70 ng RNA in 96.8 μl water, 2 μl 1M Tris•HCl (pH 7.4, Carl Roth), 0.2 μl 0.5M EDTA (Carl Roth), 1 μl 10 mg/ml MTSEA-XX-Biotin (Biotium). The reaction was incubated at room temperature for 30 to 60 min in the dark. We then separated the biotinylated RNA from the excess biotin by adding the same volume of Phenol:Chloroform:Isoamylalkohol (Sigma-Aldrich), mixing well and spinning in Phase-Lock-Gel tubes (Quantabio) at 15,000 g for 5 min. The RNA was then transferred on a Hyperbond N+ membrane (Amersham) and UV crosslinked with 2,400 μJ (254nm). To block nonspecific signal, we incubated the membrane in blocking solution (PBS pH 7.5 (Gibco), 10% SDS (Roti®-Stock 20 % SDS, Carl Roth), 1 mM EDTA) for 30 min. The membrane was then probed with a 1:5,000 dilution of 1 mg/mL streptavidin-horseradish peroxidase (Pierce) in blocking solution for 15 min. Finally the membrane was washed six times in PBS containing decreasing concentrations of SDS (10%, 1%, and 0.1% SDS, applied twice each) for 10 min. The signal of biotin-bound HRP was visualized using Amersham ECL Western Blotting Detection Reagent (GE Healthcare).

Flp-In™ 293 cells (Thermo Fisher) used as a positive and negative control were grown in DMEM (Gibco) + 10% FBS (Gibco) + 2mM L-Glutamine (Gibco) at 37°C and 5% CO_2_. The cells were incubated with 300 μM 4sU or mock treated for 15 minutes before we fixed them in methanol as described above.

### Quantification and statistical analysis

#### Mapping of tomo-seq data

Fastq files were mapped with STAR (v2.5.3a) using the –quantMode option. Genome versions used were GRCz11.95 (D. rerio), 9.2 (X. laevis) and 9.1 (X. tropicalis). From the SAM file, gene counts were assigned to a spatial barcode resulting in a count matrix.

#### Further processing of tomo-seq data

We filtered out sections with a low recovery of ERCC spike-in controls. The cutoff depends on the sequencing depth, and was set as ~0.04 percent of the mapped reads of a library (or 8000 transcripts for the replicate shown in Figure 1). In the remaining sections, we excluded lowly expressed genes (with less than 5 counts in at least one section, for the replicate shown in Figure), then divided gene counts by total counts in that section and normalized to the median section size. For clustering based on *self-organizing maps* (SOM), we calculated cumulative expression going from low to high section numbers, normalized the maximum of the cumulative expression to one and let the SOM sort these patterns into a linear matrix of 1×50 profiles. A gene was called vegetally localized in all replicates when it was assigned any profile between 46 and 50 in all replicates and at least 48 in one replicate.

#### Isoform analysis and kmer enrichment

Isoform expression in zebrafish one-cell stage embryos was determined using cufflinks v2.2.1. For kmer enrichment, we extracted 3’ UTR sequences as annotated in the zebrafish genome version GRCz10. Next, we compared vegetally localized to all expressed genes with DREME (v4.11.2) using the parameters: -g 1000 -norc -e 0.5 -mink 3 -maxk 10. For xenopus, we used the longest annotated 3’UTR for our analysis.

We calculated the 3’UTR length of a gene ID as shown in Fig. 2a by weighing the isoforms 3’UTR length according to their relative contribution to a gene IDs total expression. CDS length as shown in Fig. 2c were calculated accordingly.

#### Alignment of UTRs from *D. rerio*,*X. laevis* and *X. tropicalis*

UTR sequences were aligned using the mafft online tool (http://mafft.cbrb.jp/alignment/server/) using the following parameters: %mafft –reorder –anysymbol –maxiterate 1000 –retree 1–genafpair input

#### scSLAM-seq mapping and analysis

Raw data was demultiplexed with cellranger mkfastq (v3.0.2), and mapped with the default parameters of cellranger (10x Genomics) to the zebrafish genome, version GRCz11.95. We used the inbuilt cell detection algorithm to create a ‘whitelist’ with all barcodes that contain cells and extracted these barcodes from the BAM file to only consist of reads from real cells. We further separated the reads in that file into labeled reads (*>* 1 T to C mutation per UMI, base quality *>*20) and unlabeled reads. We then created a fastq file for labeled and for unlabeled reads, respectively, mapped them with STARsolo and obtained count matrices that were further analysed with Seurat v.3.1.2.

#### Calculation of false negative rate in scSLAM-seq

We estimated the false negative rate (i.e. the probability of a zygotic transcript molecule to remain unlabeled) with the following back-of-the-envelope calculation: We expect that approximately 5% of all Us are labeled in a zygotic transcript (Fig. 3c). The read length was 99 nt. Since the library was sequenced with ~4 reads per UMI, we assume an effective read length of 300 nt, taking into account that different reads for the same UMI may partially overlap. The GC content is on average 40%, which results in 30% Us, and hence 90 Us per transcript molecule. The probability that a zygotic transcript does not contain a single labelled U is therefore 0.95^90^ ≈ 1%.

## Supplementary tables

Supplementary Table 1: Expressed zebrafish genes and SOM profiles for three replicates

Supplementary Table 2: Marker genes for cell identities, based on labeled RNA, zebrafish, 6 hpf

Supplementary Table 3: Marker genes for cell identities, based on unlabeled RNA, zebrafish, 6 hpf

Supplementary Table 4: Expressed genes in X. laevis and SOM profiles for two replicates

Supplementary Table 5: Expressed genes in X. tropicalis and SOM profiles for two replicates

## References

1. Hashimshony, T. et al. CEL-Seq2: sensitive highly-multiplexed single-cell RNA-Seq. Genome Biol. 17, 77 (2016).

2. Islam, S. et al. Quantitative single-cell RNA-seq with unique molecular identifiers. Nat. Methods 11, 163–166 (2014).

3. Klein, A. M. et al. Droplet barcoding for single-cell transcriptomics applied to embryonic stem cells. Cell 161, 1187–1201 (2015).

4. Macosko, E. Z. et al. Highly Parallel Genome-wide Expression Profiling of Individual Cells Using Nanoliter Droplets. Cell 161, 1202–1214 (2015).

5. Haghverdi, L., Büttner, M., Wolf, F. A., Buettner, F. & Theis, F. J. Diffusion pseudotime robustly reconstructs lineage branching. Nat. Methods 13, 845–848 (2016).

6. Kester, L. & van Oudenaarden, A. Single-Cell Transcriptomics Meets Lineage Tracing. Cell Stem Cell 23 166–179 (2018).

7. Setty, M. et al. Wishbone identifies bifurcating developmental trajectories from single-cell data. Nature Biotechnology 34 637–645 (2016).

8. Trapnell, C. et al. The dynamics and regulators of cell fate decisions are revealed by pseudotemporal ordering of single cells. Nat. Biotechnol. 32, 381–386 (2014).

9. Wagner, D. E. et al. Single-cell mapping of gene expression landscapes and lineage in the zebrafish embryo. Science 360, 981–987 (2018).

10. Farrell, J. A. et al. Single-cell reconstruction of developmental trajectories during zebrafish embryogenesis. Science 360, (2018).

11. Cao, J. et al. The single-cell transcriptional landscape of mammalian organogenesis. Nature 566, 496–502 (2019).

12. Plass, M. et al. Cell type atlas and lineage tree of a whole complex animal by single-cell transcriptomics. Science 360, (2018).

13. Escobar-Aguirre, M., Elkouby, Y. M. & Mullins, M. C. Localization in Oogenesis of Maternal Regulators of Embryonic Development. Advances in Experimental Medicine and Biology 173–207 (2017) doi:10.1007/978-3-319-46095-6_5.

14. Elkouby, Y. M. & Mullins, M. C. Coordination of cellular differentiation, polarity, mitosis and meiosis – New findings from early vertebrate oogenesis. Developmental Biology 430 275–287 (2017).

15. Lein, E., Borm, L. E. & Linnarsson, S. The promise of spatial transcriptomics for neuroscience in the era of molecular cell typing. Science 358, 64–69 (2017).

16. Moor, A. E. & Itzkovitz, S. Spatial transcriptomics: paving the way for tissue-level systems biology. Curr. Opin. Biotechnol. 46, 126–133 (2017).

17. Rodriques, S. G. et al. Slide-seq: A scalable technology for measuring genome-wide expression at high spatial resolution. Science 363, 1463–1467 (2019).

18. Eng, C.-H. L. et al. Transcriptome-scale super-resolved imaging in tissues by RNA seqFISH. Nature 568, 235–239 (2019).

19. Xia, C., Fan, J., Emanuel, G., Hao, J. & Zhuang, X. Spatial transcriptome profiling by MERFISH reveals subcellular RNA compartmentalization and cell cycle-dependent gene expression. Proc. Natl. Acad. Sci. U. S. A. 116, 19490–19499 (2019).

20. Codeluppi, S. et al. Spatial organization of the somatosensory cortex revealed by osmFISH. Nat. Methods 15, 932–935 (2018).

21. Padrón, A., Iwasaki, S. & Ingolia, N. T. Proximity RNA Labeling by APEX-Seq Reveals the Organization of Translation Initiation Complexes and Repressive RNA Granules. Mol. Cell 75, 875–887.e5 (2019).

22. Fazal, F. M. et al. Atlas of Subcellular RNA Localization Revealed by APEX-Seq. Cell 178, 473–490.e26 (2019).

23. Coleman, R. A. et al. Imaging Transcription: Past, Present, and Future. Cold Spring Harb. Symp. Quant. Biol. 80, 1–8 (2015).

24. La Manno, G. et al. RNA velocity of single cells. Nature 560, 494–498 (2018).

25. Erhard, F. et al. scSLAM-seq reveals core features of transcription dynamics in single cells. Nature 571, 419–423 (2019).

26. Hendriks, G.-J. et al. NASC-seq monitors RNA synthesis in single cells. Nat. Commun. 10, 3138 (2019).

27. Battich, N. et al. Sequencing metabolically labeled transcripts in single cells reveals mRNA turnover strategies. Science 367, 1151–1156 (2020).

28. Cao, J., Zhou, W., Steemers, F. et al. Sci-fate characterizes the dynamics of gene expression in single cells. Nat Biotechnol 38, 980–988 (2020).

29. Qiu, Q. et al. Massively parallel and time-resolved RNA sequencing in single cells with scNT-seq. Nat. Methods 17, 991–1001 (2020).

30. Junker, J. P. et al. Genome-wide RNA Tomography in the zebrafish embryo. Cell 159, 662–675 (2014).

31. Holler, K. & Junker, J. P. RNA Tomography for Spatially Resolved Transcriptomics (Tomo-Seq). Methods Mol. Biol. 1920, 129–141 (2019).

32. Kohonen, T. Self-organized formation of topologically correct feature maps. Biological Cybernetics 43 59–69 (1982).

33. Kimmel, C. B., Ballard, W. W., Kimmel, S. R., Ullmann, B. & Schilling, T. F. Stages of embryonic development of the zebrafish. Dev. Dyn. 203, 253–310 (1995).

34. Langdon, Y. G. & Mullins, M. C. Maternal and zygotic control of zebrafish dorsoventral axial patterning. Annu. Rev. Genet. 45, 357–377 (2011).

35. Welch, E. & Pelegri, F. Cortical depth and differential transport of vegetally localized dorsal and germ line determinants in the zebrafish embryo. Bioarchitecture 5, 13–26 (2014).

36. Schier, A. F. & Talbot, W. S. Molecular genetics of axis formation in zebrafish. Annu. Rev. Genet. 39, 561–613 (2005).

37. Herzog, V. A. et al. Thiol-linked alkylation of RNA to assess expression dynamics. Nat. Methods 14, 1198–1204 (2017).

38. Cao, J., Zhou, W., Steemers, F., Trapnell, C. & Shendure, J. Sci-fate characterizes the dynamics of gene expression in single cells. Nat. Biotechnol. 38, 980–988 (2020).

39. Heyn, P. et al. The earliest transcribed zygotic genes are short, newly evolved, and different across species. Cell Rep. 6, 285–292 (2014).

40. Yoon, C., Kawakami, K. & Hopkins, N. Zebrafish vasa homologue RNA is localized to the cleavage planes of 2- and 4-cell-stage embryos and is expressed in the primordial germ cells. Development 124, 3157–3165 (1997).

41. Raz, E. Primordial germ-cell development: the zebrafish perspective. Nat. Rev. Genet. 4, 690–700 (2003).

42. Knaut, H., Steinbeisser, H., Schwarz, H. & Nüsslein-Volhard, C. An evolutionary conserved region in the vasa 3’UTR targets RNA translation to the germ cells in the zebrafish. Curr. Biol. 12, 454–466 (2002).

43. Claussen, M., Horvay, K. & Pieler, T. Evidence for overlapping, but not identical, protein machineries operating in vegetal RNA localization along early and late pathways in Xenopus oocytes. Development 131, 4263–4273 (2004).

44. Claußen, M. et al. Global analysis of asymmetric RNA enrichment in oocytes reveals low conservation between closely related Xenopus species. Mol. Biol. Cell 26, 3777–3787 (2015).

45. Owens, D. A. et al. High-throughput analysis reveals novel maternal germline RNAs crucial for primordial germ cell preservation and proper migration. Development 144, 292–304 (2017).

46. Sindelka, R. et al. Asymmetric distribution of biomolecules of maternal origin in the Xenopus laevis egg and their impact on the developmental plan. Sci. Rep. 8, 8315 (2018).

47. Rothschild, S. C. et al. CaMK-II activation is essential for zebrafish inner ear development and acts through Delta-Notch signaling. Dev. Biol. 381, 179–188 (2013).

48. Betley, J. N., Frith, M. C., Graber, J. H., Choo, S. & Deshler, J. O. A ubiquitous and conserved signal for RNA localization in chordates. Curr. Biol. 12, 1756–1761 (2002).

49. Jambhekar, A. & Derisi, J. L. Cis-acting determinants of asymmetric, cytoplasmic RNA transport. RNA 13, 625–642 (2007).

50. Kosaka, K., Kawakami, K., Sakamoto, H. & Inoue, K. Spatiotemporal localization of germ plasm RNAs during zebrafish oogenesis. Mech. Dev. 124, 279–289 (2007).

51. Taliaferro, J. M. et al. Distal Alternative Last Exons Localize mRNAs to Neural Projections. Mol. Cell 61, 821–833 (2016).

52. Trapnell, C. et al. Transcript assembly and quantification by RNA-Seq reveals unannotated transcripts and isoform switching during cell differentiation. Nat. Biotechnol. 28, 511–515 (2010).

53. Yartseva, V., Takacs, C. M., Vejnar, C. E., Lee, M. T. & Giraldez, A. J. RESA identifies mRNA-regulatory sequences at high resolution. Nat. Methods 14, 201–207 (2017).

54. Bailey, T. L. DREME: motif discovery in transcription factor ChIP-seq data. Bioinformatics 27, 1653–1659 (2011).

55. Rabani, M., Pieper, L., Chew, G.-L. & Schier, A. F. A Massively Parallel Reporter Assay of 3’ UTR Sequences Identifies In Vivo Rules for mRNA Degradation. Mol. Cell 68, 1083–1094.e5 (2017).

56. Chang, P. et al. Localization of RNAs to the mitochondrial cloud in Xenopus oocytes through entrapment and association with endoplasmic reticulum. Mol. Biol. Cell 15, 4669–4681 (2004).

57. King, M. L., Messitt, T. J. & Mowry, K. L. Putting RNAs in the right place at the right time: RNA localization in the frog oocyte. Biol. Cell 97, 19–33 (2005).

58. Jambor, H. et al. Systematic imaging reveals features and changing localization of mRNAs in Drosophila development. Elife 4, (2015).

59. Lécuyer, E. et al. Global analysis of mRNA localization reveals a prominent role in organizing cellular architecture and function. Cell 131, 174–187 (2007).

60. Moor, A. E. et al. Global mRNA polarization regulates translation efficiency in the intestinal epithelium. Science 357, 1299–1303 (2017).

61. Ciolli Mattioli, C. et al. Alternative 3′ UTRs direct localization of functionally diverse protein isoforms in neuronal compartments. Nucleic Acids Research 47 2560–2573 (2019).

62. Zappulo, A. et al. RNA localization is a key determinant of neurite-enriched proteome. Nat. Commun. 8, 583 (2017).

63. Rabani, M., Kertesz, M. & Segal, E. Computational prediction of RNA structural motifs involved in posttranscriptional regulatory processes. Proc. Natl. Acad. Sci. U. S. A. 105, 14885–14890 (2008).

64. Martin, K. C. & Ephrussi, A. mRNA localization: gene expression in the spatial dimension. Cell 136, 719–730 (2009).

65. Vourekas, A., Alexiou, P., Vrettos, N., Maragkakis, M. & Mourelatos, Z. Sequence-dependent but not sequence-specific piRNA adhesion traps mRNAs to the germ plasm. Nature 531, 390–394 (2016).

66. Grün, D., Kester, L. & van Oudenaarden, A. Validation of noise models for single-cell transcriptomics. Nat. Methods 11, 637–640 (2014).

67. Claussen, M. & Pieler, T. Identification of vegetal RNA-localization elements in Xenopus oocytes. Methods 51, 146–151 (2010).

68. Thisse, C. & Thisse, B. High-resolution in situ hybridization to whole-mount zebrafish embryos. Nat. Protoc. 3, 59–69 (2008).

