## Supplemental_Figures for "Spatio-temporal mRNA dynamics in the early zebrafish embryo"

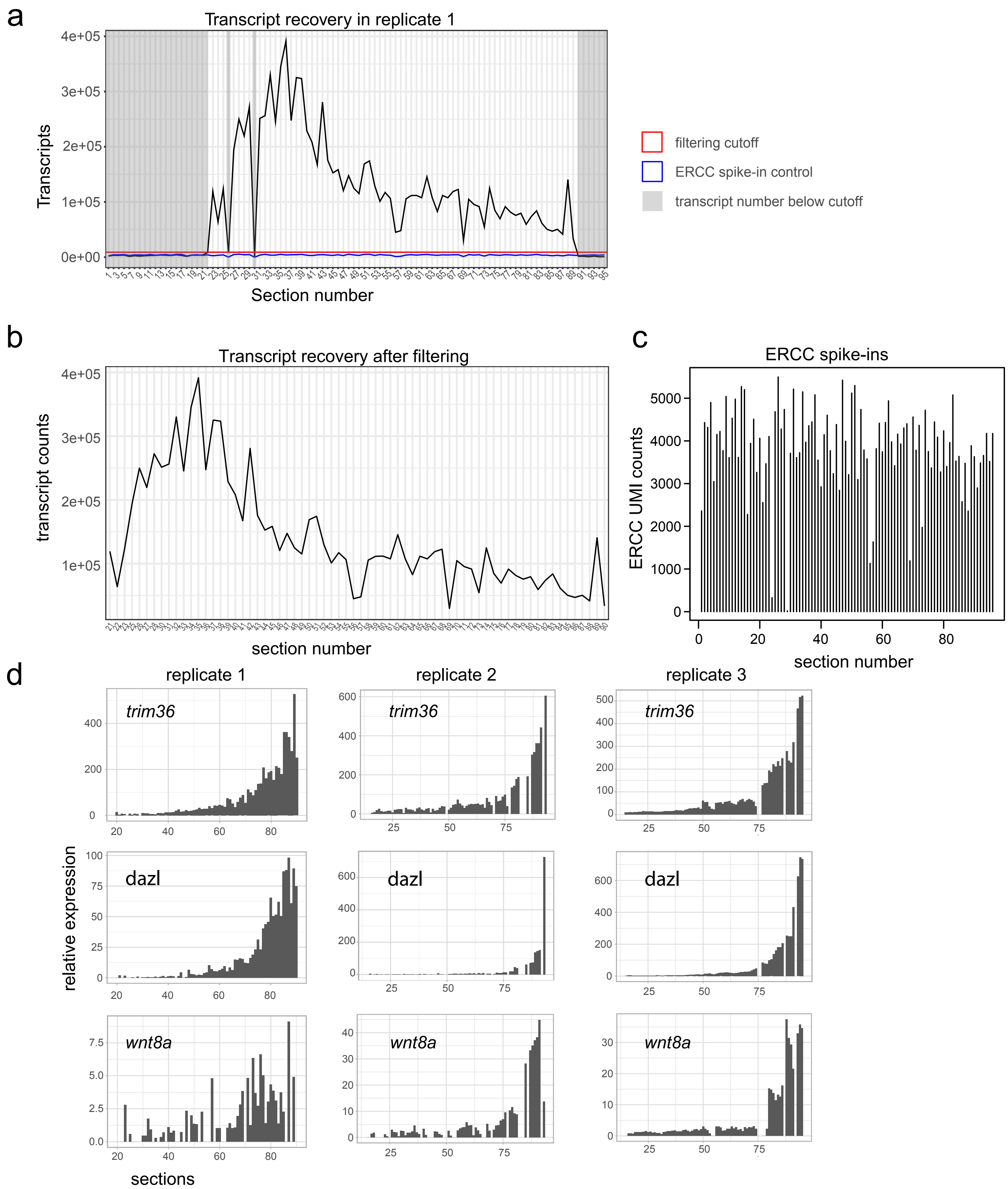


**Supplemental Figure 1: Tomo-seq data processing.** (a). Summed up transcript recovery per section for zebrafish tomo-seq replicate 1. The embryo was cut along the animal-vegetal axis. Filtering cut-off for unsuccessful sections in red, the sections that have transcript sums below that cut-off are shaded in gray. The blue line indicates counts for ERCC spike-in molecules. (b). The transcript profile from a, after the filtering step and before normalization.
(c). ERCC spike-in UMI counts per section for zebrafish replicate 1.
(d). Tomo-seq tracks for known vegetally localizing genes trim36, dazl, wnt8a in all three zebrafish tomo-seq replicates.

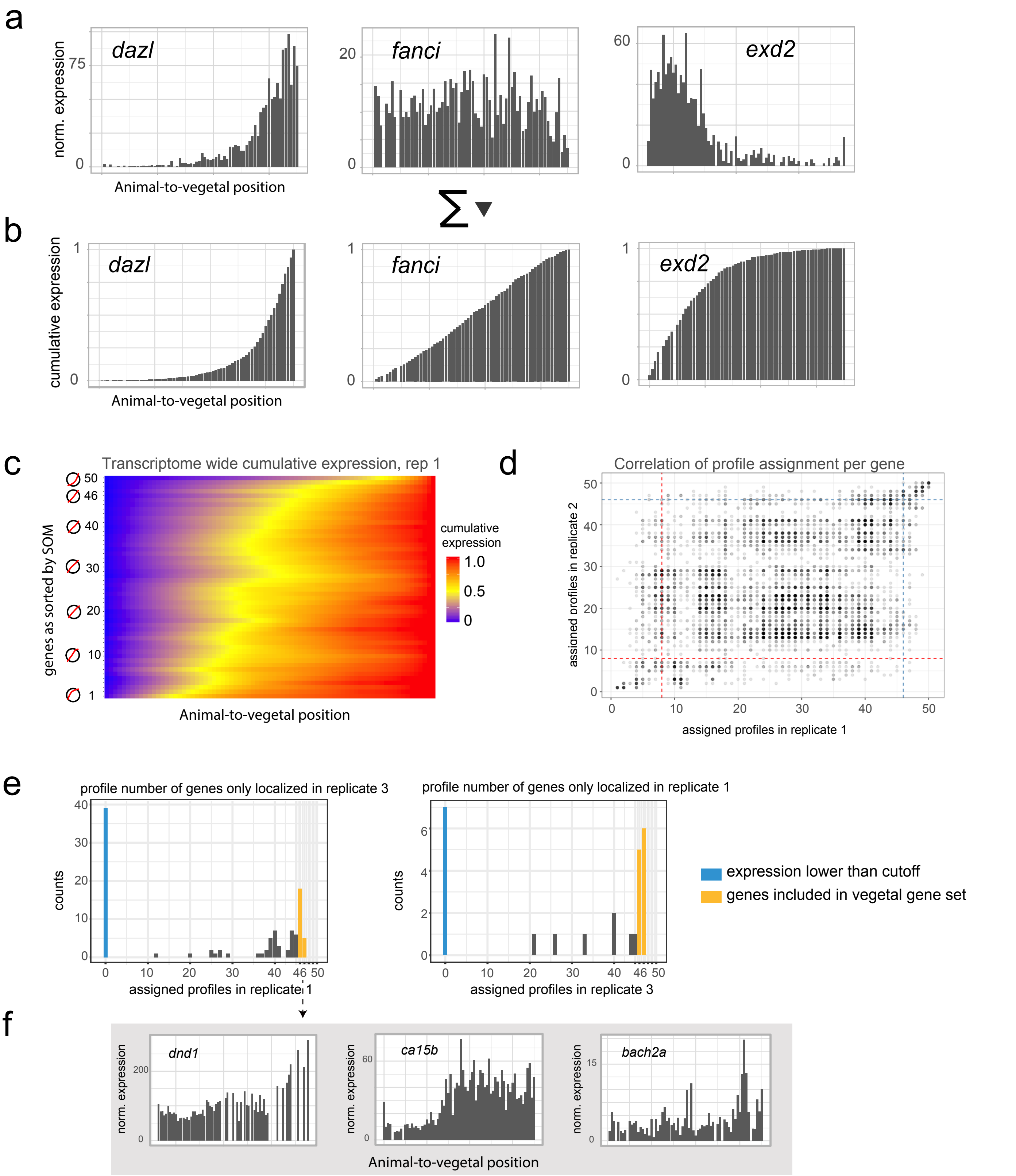


**Supplemental Figure 2: Zebrafish tomo-seq data analysis.**

(a) Exemplary tomo-seq tracks for genes that are either vegetally localized, not localized or animally localized.

(b) Cumulative expression patterns for the genes in A, with the sum of counts for a specific gene normalized to 1.

(c) Heatmap of cumulative expression patterns. Spatial position in the embryo on the x-axis, genes as sorted by SOM into profiles 1-50 on the y-axis. Animally localized genes are at the bottom.

(d) Correlation of profile assignment to all genes from two replicates. Profiles 1 to 50 of replicate 1 on the x-axis, profiles 1-50 of replicate 2 on the y-axis; dots are genes. Dashed lines are cutoffs for animal and vegetal localization, respectively.

(e) Histogram of profile numbers of genes that are not localized in both of the replicates 1 and 3. Most of the genes were only recovered in one sample (blue bar), or fall into profiles just below the cutoff (orange). Left side: Profiles of genes assigned vegetal localization replicate 3, but not in replicate 1. Right side: Profiles of genes of replicate 3, found vegetally localized only in replicate 1.

(f) Exemplary tomo-seq tracks of genes *dnd1*, *ca15b* and *bach2a* from the orange bars in e.

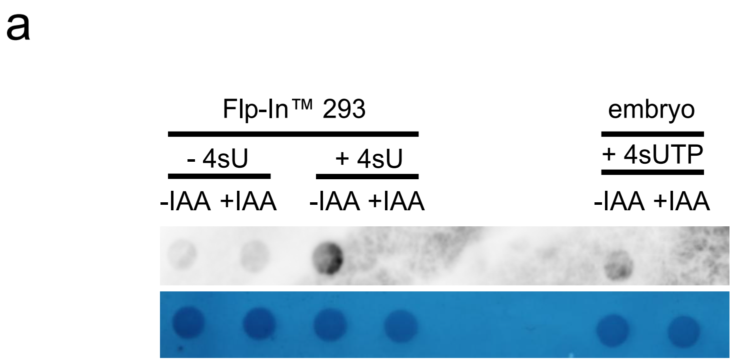


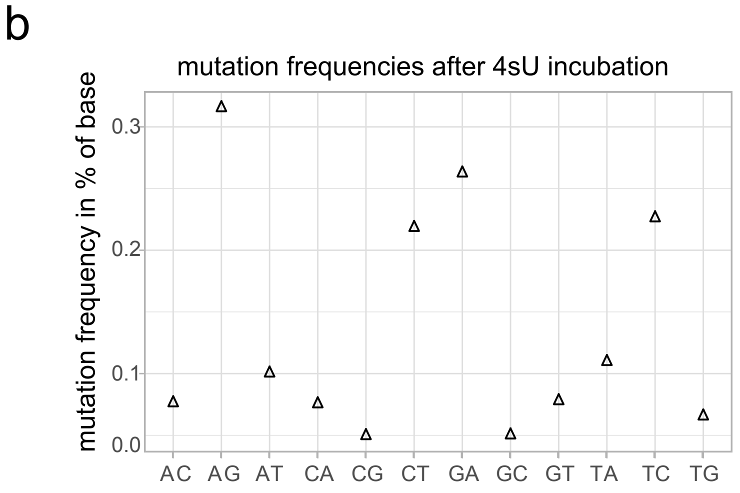


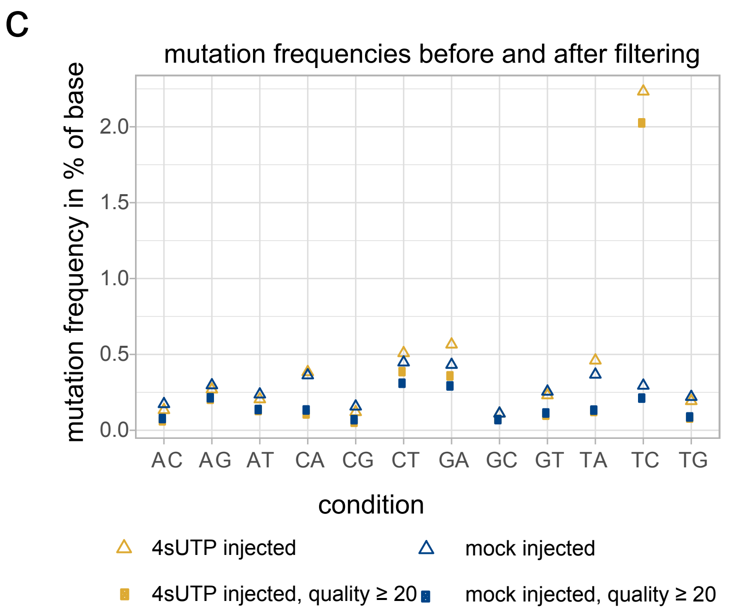


**Supplemental Figure 3: additional data for the scSLAMseq method:**

(a). Label incorporation and IAA derivatization in fixed cells. Flp-In™ 293 cells treated with 300µM 4sU or mock treated for 15 min and zebrafish embryos injected with 4nl 12.5mM 4sUTP. IAA conversion was performed in methanol-fixed cells (Methods). Disappearing signal indicates successful derivatization.
(b). Base mutation frequencies of a bulk library obtained by incubating dechorionated zebrafish embryos in 100mM 4sU for 90 minutes prior to collection at shield stage. Sequenced on a NextSeq500 machine (Illumina).
(c). Base mutation frequencies of a scSLAM-seq library before and after quality filtering, sequenced on a NextSeq500 machine (Illumina). Reads with at least one labeling event in yellow, unlabeled reads in blue.

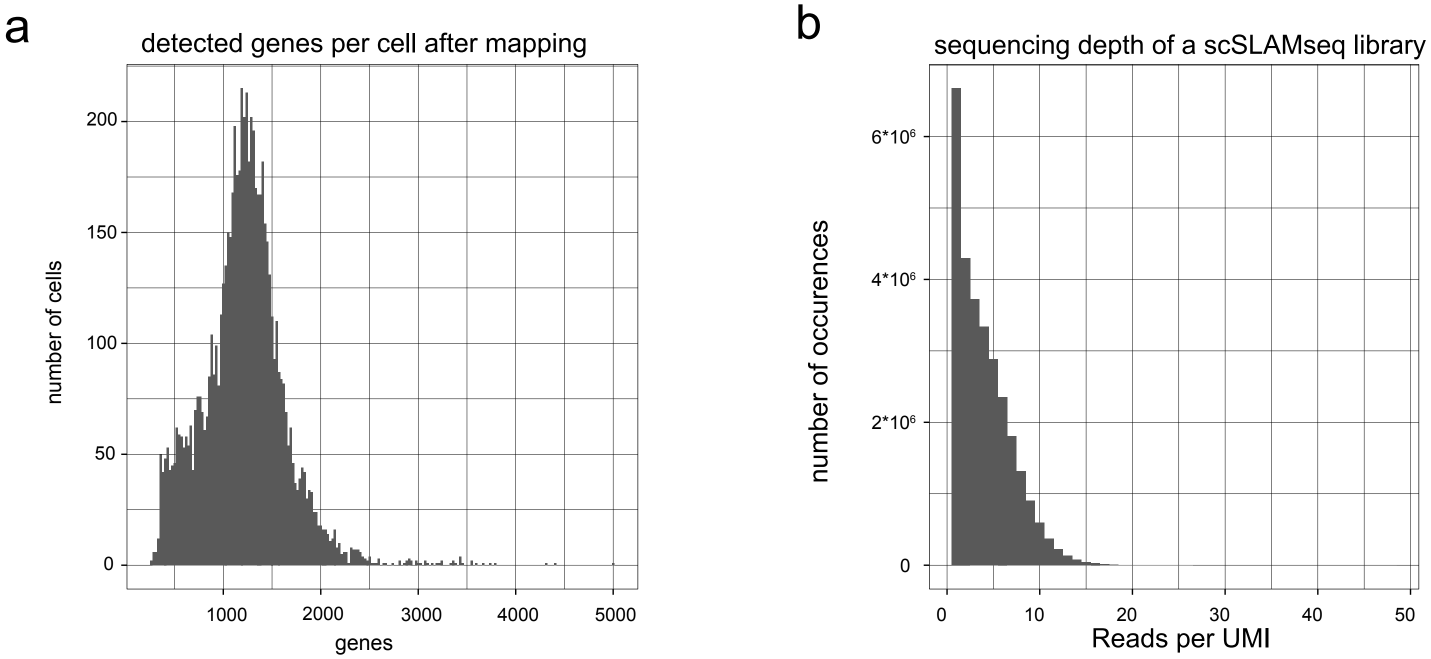


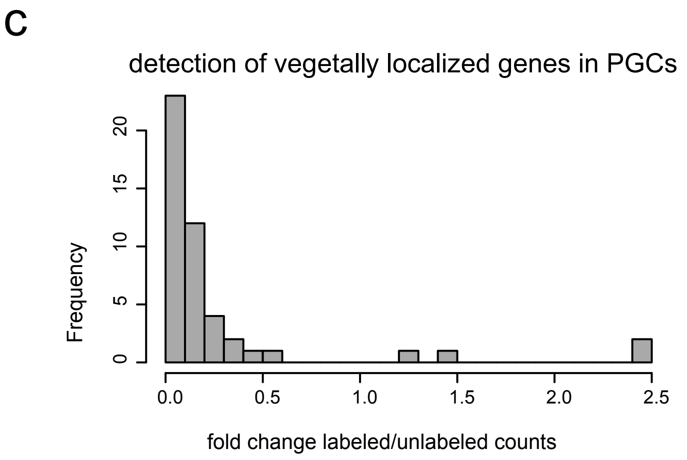


**Supplemental Figure 4: Maternal vegetally localized genes at gastrulation stage:**

(a) Histogram of total number of detected genes per cell after mapping, median is 1218.
(b) Sequencing depth of the scSLAM-seq librar y, determined as ‘UMI coverage’, or reads per UMI (mean = 4.07).
(c) Fold change of labeled to unlabeled counts for maternal vegetally localized genes in PGCs. Most of these genes are barely expressed zygotically by primordial germ cells.

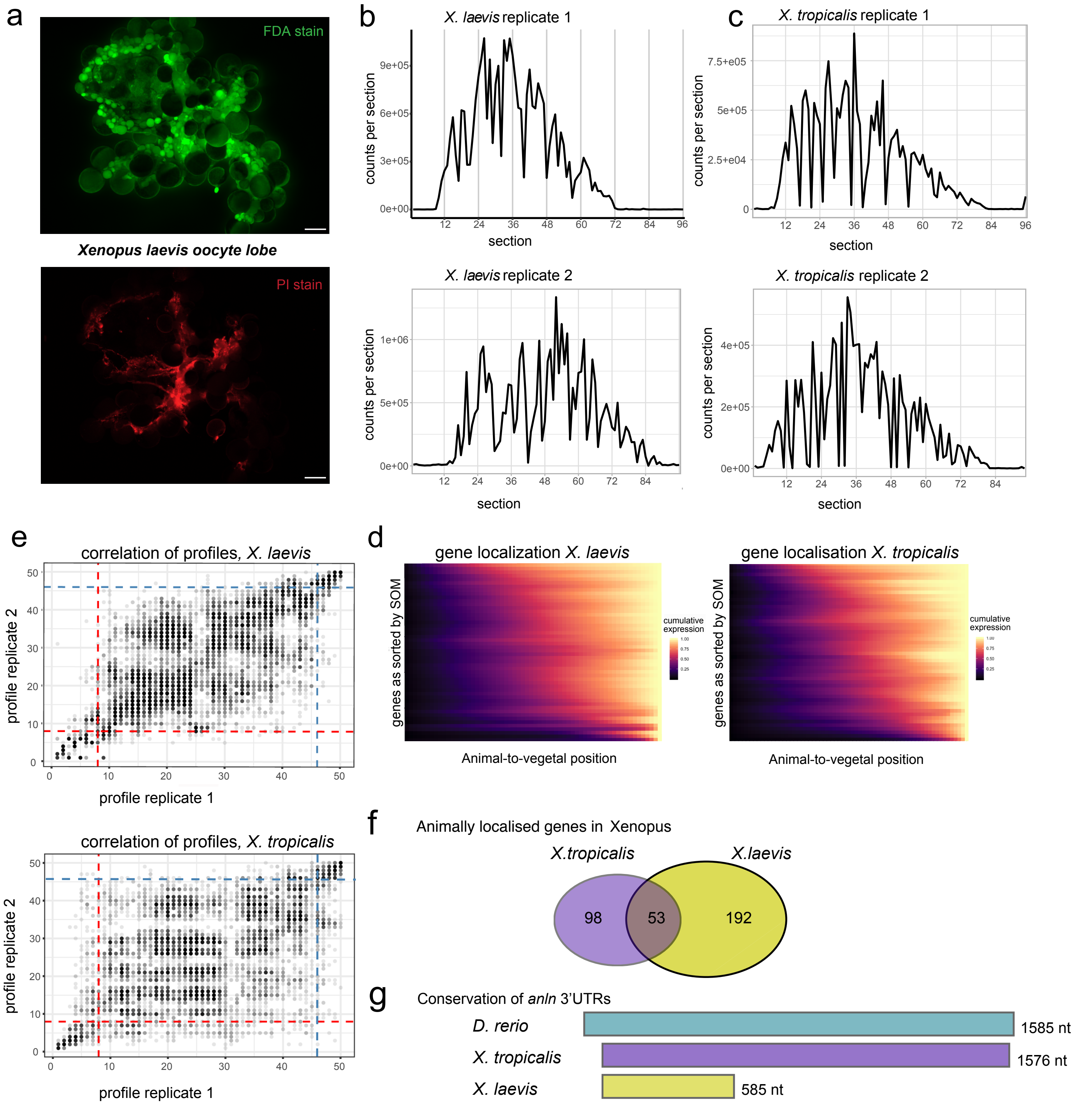


**Supplemental Figure 5: Additional data for xenopus tomo-seq.**

(a). Live (FDA) and dead (PI) stain for xenopus oocytes upon arrival, prior to manual and enzymatic dissection. Scale bars represent 1 mm.

(b) and (c). Total transcript counts for two tomo-seq replicates from (b) *X. laevis* and (c) *X. tropicalis*.

(d). Heatmap of cumulative expression patterns for xenopus species. Spatial position in the embryo on the x-axis, genes as sorted by SOM into profiles 1-50 on the y-axis. Animally localized genes are at the top.

(e). Correlation of profile assignment to all genes of two replicates from *X. laevis* and *X. tropicalis*. Profiles 1-50 of replicate 1 on the x-axis, profiles 1-50 of replicate 2 on the y-axis; dots are genes. Dashed lines are cutoffs for animal and vegetal localization, respectively.

(f). Overlap of animally localized genes in xenopus species. Only genes that were detected in both species were considered.

(g). Alignment of *anln* 3'UTRs from *D. rerio*, *X. laevis* and *X. tropicalis*. Xenopus sequences were obtained from xenbase, for zebrafish the highest expressed 3'UTR isoform was used. Sequences were aligned with mafft (see Methods).

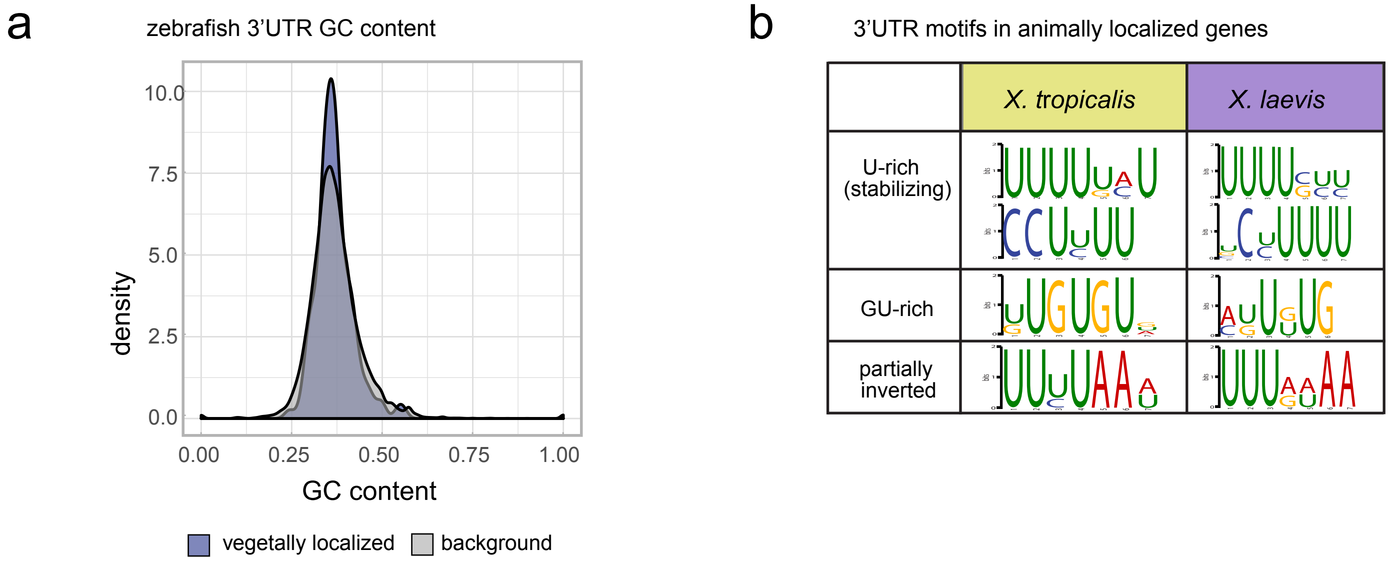


**Supplemental Figure 6: 3’UTR characteristics and kmer enrichment analysis.**

(a). Comparison of sequence characteristics of 216 expressed isoforms (Trapnell *et al.*, 2010) of vegetally localized to all genes: GC content, mean = 0.37.
(b). Results of the kmer enrichment analysis of the longest 3’UTR of animally localized genes in *X. laevis* and *X. tropicalis*, top four motifs. Resulting motif logos of both species were grouped by similarity.
